# Spatio-temporal T cell tracking for personalized TCR-T designs in childhood cancer

**DOI:** 10.1101/2024.11.18.624071

**Authors:** Inés Sentís, Juan L. Melero, Alex Cebria-Xart, Marta Grzelak, Marta Soto, Alexandra Michel, Quirze Rovira, Carlos J. Rodriguez-Hernandez, Ginevra Caratù, Andrea Urpi-Badell, Christophe Sauvage, Ana Mendizabal-Sasieta, Davide Maspero, Anna Pascual-Reguant, Juan Pablo Muñoz Perez, Jaume Mora, Alexandre Harari, Juan C. Nieto, Alexandra Avgustinova, Holger Heyn

**Author notes:** Authors contributed equally (Juan L. Melero, Alex Cebria-Xart). Corresponding authors: Alexandra Avgustinova and Holger Heyn.

## Abstract

Immune checkpoint inhibition (ICI) has revolutionized oncology, offering extended survival and long-term remission in previously incurable cancers. While highly effective in tumors with high mutational burden, lowly mutated cancers, including pediatric malignancies, present low response rate and limited predictive biomarkers. Here, we present a framework for the identification and validation of tumor-reactive T cells as a biomarker to quantify ICI efficacy and as candidates for a personalized TCR-T cell therapy. Therefore, we profiled a pediatric malignant rhabdoid tumor patient with complete remission after ICI therapy using deep single-cell T cell receptor (TCR) repertoire sequencing of the tumor microenvironment (TME) and the peripheral blood. Specifically, we tracked T cell dynamics longitudinally from the tumor to cells in circulating over a time course of 12 months, revealing a systemic response and durable clonal expansion of tumor-resident and ICI-induced TCR clonotypes. We functionally validated tumor reactivity of TCRs identified from the TME and the blood by co-culturing patient-derived tumor cells with TCR-engineered autologous T cells. Here, we observed unexpectedly high frequencies of tumor-reactive TCR clonotypes in the TME and confirmed T cell dynamics in the blood post-ICI to predict tumor-reactivity. These findings strongly support spatio-temporal tracking of T cell activity in response to ICI to inform therapy efficacy and to serve as a source of tumor-reactive TCRs for personalized TCR-T designs.

## Main

Immune checkpoint inhibition (ICI) therapy has shown tremendous efficacy in cancer types, such as cutaneous melanoma^1^, colorectal cancer^2,3^ (CRC), and non-small cell lung cancer^4–6^ (NSCLC). Specifically, tumors with high mutational burden induced by external factors (e.g., UV radiation and smoke) or tumor intrinsic alterations (e.g., microsatellite instability, MSI) show high response rate, likely due to the increased presentation of neo-antigens that make the tumor visible to the immune system^7^. For low-mutational cancer types, alternative biomarkers have been suggested in both the tumor (e.g., PD-L1 expression) and the tumor microenvironment (TME; e.g., CXCL13) compartment, however, their correlation with response is generally low and novel patient monitoring, stratification and immunotherapy strategies are an unmet clinical need^7,8^. This challenge is particularly evident in childhood cancers, which typically have a low tumor mutational burden (TMB) compared to adult cancers^9^. Nevertheless, some children with cancer show objective clinical benefit from ICI, highlighting that other factors in addition to TMB must determine ICI success in these patients^10,11^. Indeed, splicing disruption or dysregulation of endogenous retroviruses have been suggested to drive immune infiltration and predict ICI response in solid childhood cancers^12,13^. However, the underlying biology that drives these alternative biomarkers is complex and, thus, more difficult to conclusively evaluate in a clinical setting.

T cell presence and, more generally, an inflamed TME has been one of the first biomarkers suggested for patient stratification in the context of ICI^14,15^. Considering that T cells are the direct effector cell type for ICI drugs, such as anti-CTLA-4 and anti-PD-1/anti-PD-L1 treatment, a T cell biomarker for response and monitoring is conceptually appealing. Accordingly, T cell activity and related T cell receptor (TCR) repertoire dynamics have been shown to be associated with ICI response and to be different between responding and non-responding patients^16^. However, predicting patient response based solely on the presence or diversity of T cells has shown only moderate success across studies and cancer types, suggesting more sophisticated T cell profiling technologies and analysis tools to be required to decipher and clinically leverage the complexity of T cell mediated anti-tumor response^7^. Particularly, the fact that tumor-reactive T cell clones are often newly induced by ICI (clonal replacement) and not just reactivated (clonal invigoration), make generalized prediction and biomarker discovery challenging^17^.

Using the TCR as a natural clonal lineage barcode, in addition to its specificity for recognizing (neo-)antigens presented on tumor cells, has immense potential in clinical diagnostics^18,19^. However, technical limitations to date impeded a widespread application despite highly promising proof-of-concept studies^20^. Technical shortcomings today include the lack of depth and accuracy when measuring T cell dynamics, which ultimately hinder the longitudinal and multi-compartment (e.g. TME and blood) clonal tracking. Here, single-cell TCR sequencing shows high accuracy for the quantification of clonotype size, and the potential to pair alpha and beta TCR chains, but it lacks the scale to comprehensively chart TCR diversity (by profiling only thousands of cells per experiment). On the other hand, bulk TCR sequencing reaches higher depths, but lacks accuracy due to technical artifacts (PCR biases) and the inability to perform receptor chain pairing. Scalability in TCR sequencing is particularly important when profiling blood samples, where single clonotypes are highly diluted and clonal frequencies are generally low. Receptor pairing is a crucial requirement to validate T cell specificity against the tumor, for which *in vitro* validation of anti-tumor reactivity is considered the gold-standard. Proxies and inference of anti-tumor reactivity, such as the T cell state (e.g., exhaustion) or antigen prediction, lack specificity to accurately pinpoint tumor reactive TCRs. Indeed, conclusive validation of tumor reactivity becomes essential when turning T cells into living drugs and the TCR into cellular therapies (e.g. TCR-T cells).

Here we report an end-to-end framework for the identification, prioritization and functional engineering of tumor-reactive TCRs in an ICI-responding childhood cancer patient. In detail, we applied a novel, deep single-cell T cell sequencing technology (OS-T) to quantify and track the response induced by ICI treatment in a pediatric malignant rhabdoid tumor (MRT) patient in the TME and longitudinally in the blood over the course of 12 months. MRTs are rare and highly aggressive tumors that primarily affect infants and young children (2 years median age of diagnosis^21^). MRTs can affect a variety of tissues, including the central nervous system (called atypical teratoid/rhabdoid tumors, ATRT) and different extracranial locations, such as kidneys (rhabdoid tumors of the kidney, RTK) and other soft tissues ^22,23^. Due to rapid and invasive primary tumor growth and limited responses to radiotherapy and chemotherapy, MRT patients have dismal prognosis, with 5-year survival rates below 20%^21^. Combined radio/chemotherapy and ICI (anti-PD-L1) treatment of the here profiled RTK patient resulted in a complete clinical response. In this context, we provide evidence for a long-lasting, ICI-driven induction of T cell activity, with therapy-expanded clonotypes detectable in the blood after 12 months. We confirmed tumor-specificity of both tumor-resident and ICI-induced clonotypes through challenge experiments with TCR-engineered autologous T cells. Overall, we validate deep single-cell TCR profiling of both TME and blood as a source for the identification of tumor-reactive T cells in low TMB childhood cancers and implement a framework for TCR engineering as a proof-of-concept for personalized cell therapy.

## Results

### Clinical course and sample collection

A patient aged 5 months was admitted with acute abdominal distension. Magnetic resonance imaging (MRI) identified the presence of a large infiltrative renal mass (**Figure 1a**), which was surgically removed by radical nephroureterectomy with retroperitoneal lymphadenectomy. The patient was subjected to an intensive chemotherapy regimen concomitant with total abdominal radiotherapy (10.5 Gy) (COG-AREN0321 protocol, **Figure 1b**). Tumor tissue biopsies were obtained from the resected lesion, along with blood samples taken at seven time points during treatment (**Figure 1b**). Whole-genome sequencing of tumor tissue revealed a low-TMB of 0.16 coding mutations per megabase^9^. Immunohistochemical staining showed the absence of INI1 (also known as SMARCB1) in the tumor cell compartment, confirming malignant rhabdoid tumor (MRT) diagnosis (**Figure 1c**). IHC for CD3 and PD-L1 identified heterogeneous T cell infiltration and PD-L1 expression, with tumor areas containing >30% PD-L1+ tumor cells, suggesting that ICI therapy may be beneficial for this patient (**Figure 1c**). Given the severity of the disease and the dismal initial prognosis, the patient received Atezolizumab (anti-PD-L1) every 21 days under compassionate use in addition to the COG-AREN0321 protocol, starting with cycle 2 of chemotherapy. Unexpectedly, by the end of the COG-AREN0321 protocol (30 weeks after tumor resection), the patient achieved complete remission (CR) based on computed tomography (CT) profiles. At this stage, both chemotherapy and atezolizumab were discontinued. Due to the positive response to treatment, the clinical decision was to continue ICI alone as maintenance therapy. However, as the patient presented multifactorial ascites and grade 3 hypertransaminasemia, likely due to immune-related hepatitis, ICI-treatment re-initiation had to be delayed while the patient was treated with corticosteroids. Once adverse symptoms subsided, and 8 weeks after discontinuation of the previous regimen, nivolumab (anti-PD1) ICI treatment was initiated at 28 day intervals as maintenance therapy. 18 months after diagnosis, the patient remains in CR, and without toxicities associated with the nivolumab treatment.

**Figure 1:**
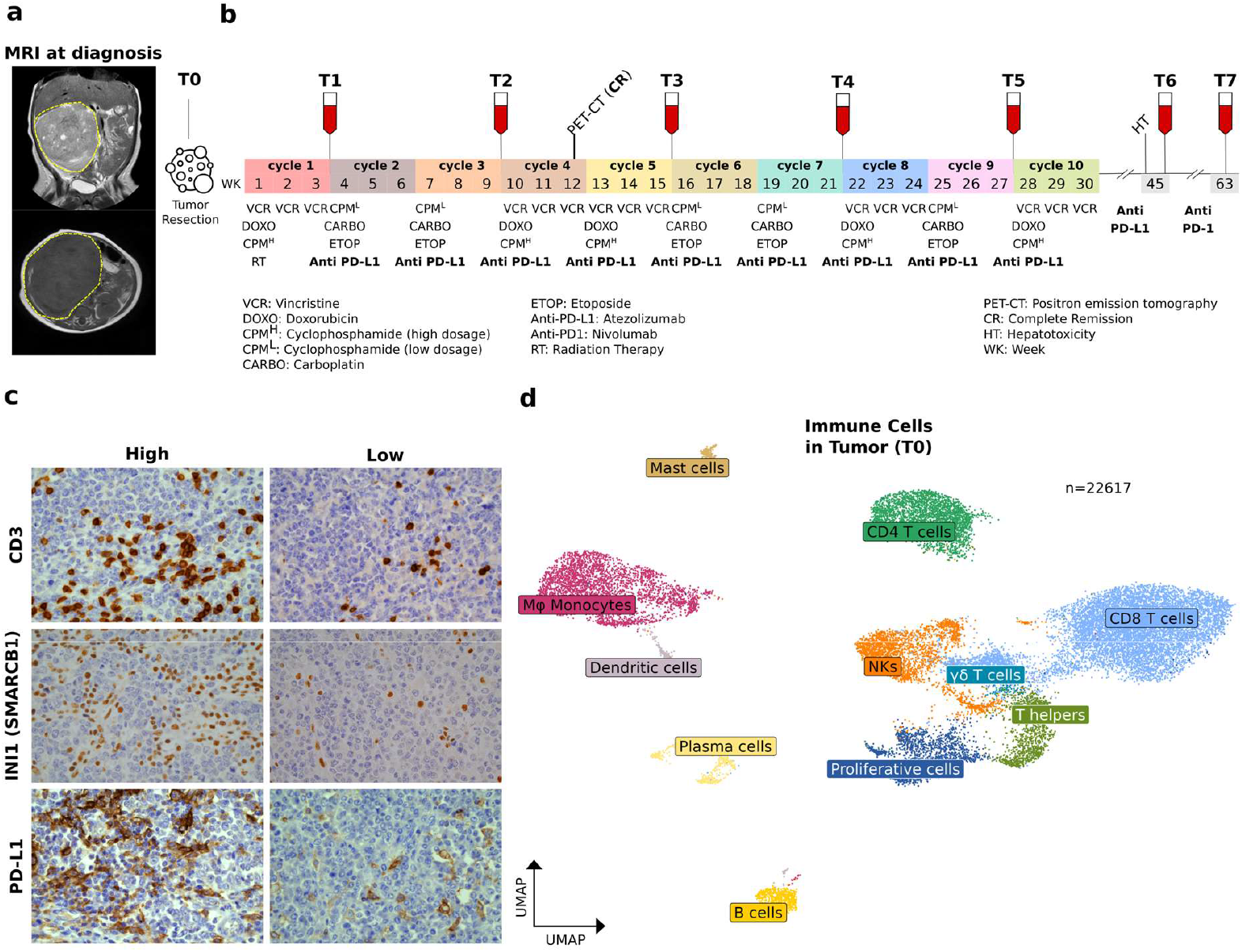
Clinical course and tumor immune profiles. (**a**) Magnetic resonance imaging (MRI) at patient diagnosis. The yellow lines delineate the primary tumor of the kidney. (**b**) Schematic timeline of the chemo/radio therapy and ICI treatment of the patient, including sample collection time points of tumor and blood specimens for this study. (**c**) Immunohistochemistry staining with CD3, INI1 (corresponding to SMARCB1 gene) and PD-L1 of the primary tumor sample. Displayed are tumor areas with high (left) and low (right) staining indicating different characteristics of immunogenicity. Scale bar (50 µm). (**d**) Uniform Manifold Approximation and Projection (UMAP) of the immune cell fraction in the TME of the primary tumor (T0), annotated with canonical gene expression markers used for cell types and states.

Considering the efficacy of the combined chemo- and immunotherapy and the potential driving role of T cells for the success of the treatment, we generated a spatio-temporal map of T cell activity of the patient. Such detailed charting of T cells served to identify, quantify and monitor both intrinsic and therapy-induced, anti-tumor immunity. Importantly, we utilized the generated information to guide informed clinical decision-making to continue ICI treatment of the patient without the combination with chemo/radiotherapy (after cycle 10) and to simulate a second-line TCR-based personalized cellular therapy.

### T cell profiles indicate an active response to the tumor

To obtain a high-resolution picture of the patient’s TME immune composition, cryopreserved tumor tissue underwent single-cell digestion and FACS sorting to isolate immune cells (CD45+), also specifically enriching for T cells (CD3+, **Supplementary Table 1**). We then performed single-cell RNA and TCR sequencing, yielding high-quality transcriptome profiles and TCR repertoires for 22,617 cells. We first annotated principal cell lineages to assign major cell types (**Figure 1d**; **Extended Data Fig.1**; **Supplementary Table 2 and 3**), before zooming into the T cell compartment with 14,789 tumor-infiltrating T cells **Figure 2a**). Major T cell lineages were represented by CD8 T cells (60%), CD4 T cells (Tregs; 20%) and proliferating cells (9%) from both CD4 and CD8 subtypes (**Figure 2b, Supplementary Table 4**). NKT cells, y/d T cells and T helper CD4 cells together represented 14% of T cells in the TME. Interestingly, most CD8 cells expressed genes indicative of cytotoxicity and markers for (pre-)exhaustion (c0 and c3, **Figure 2c, Supplementary Table 4**), indicating their prior activity within the tumor. Another prominent CD8 cluster (c10) presented high expression of CXCL13 and TNFRSF9 (gene encoding for 4-1BB protein), known markers for activation and reactivity of tumor-infiltrating T cells and associated with response to ICI^24^. In addition, we detected CD8 tumor-resident memory (RM) cells with high expression of *ZNF683* (known as Hobit protein; c4), a known marker for tissue resident T cells^25^. Analysis of TCR clonotype expansion within the TME, revealed a strong clonal amplification of IFNG+ and (pre-)exhausted CD8 T cells (c0, c1, c3), but also showed expansion of Treg cells (c2, c5), and to a lower extent of T helper CD4 cells (c6, **Figure 2d,e**). Within the CD4 compartment, we identified three different Treg populations (c2,c5 and c13) in addition to proliferating Treg cells (c9_1; **Figure 2f, Extended Data Fig.2, Supplementary Table 4**). In addition, non-Treg CD4 cells expressed high levels of CD40LG, indicating a T helper phenotype (c9_0, c6, c11; **Extended Data Fig.2**). We re-clustered CD4 T helper populations to gain granularity of different functional subtypes, identifying Th1, Th2, Th17 and T follicular helper cells (Tfh, **Figure 2g**,**h, Supplementary Table 5**). Overall, the presence of clonally amplified (pre-)exhausted CD8 cells and the ongoing proliferation of T cells pointed to a highly inflamed TME. Conversely, the high presence of Treg populations within the tumor suggests an ongoing arms-race between the inflammatory, anti-tumor CD8 T cell response, and immunosuppressive, tumor-promoting Treg populations, further highlighting ICI intervention as a promising clinical course of action. Moreover, the strong clonal and phenotypic diversity of tumor-resident T helper cells, together with the presence of B and Plasma cells provided the basis for building a long-lasting adaptive anti-tumor immunity (**Figure 1d**).

**Figure 2:**
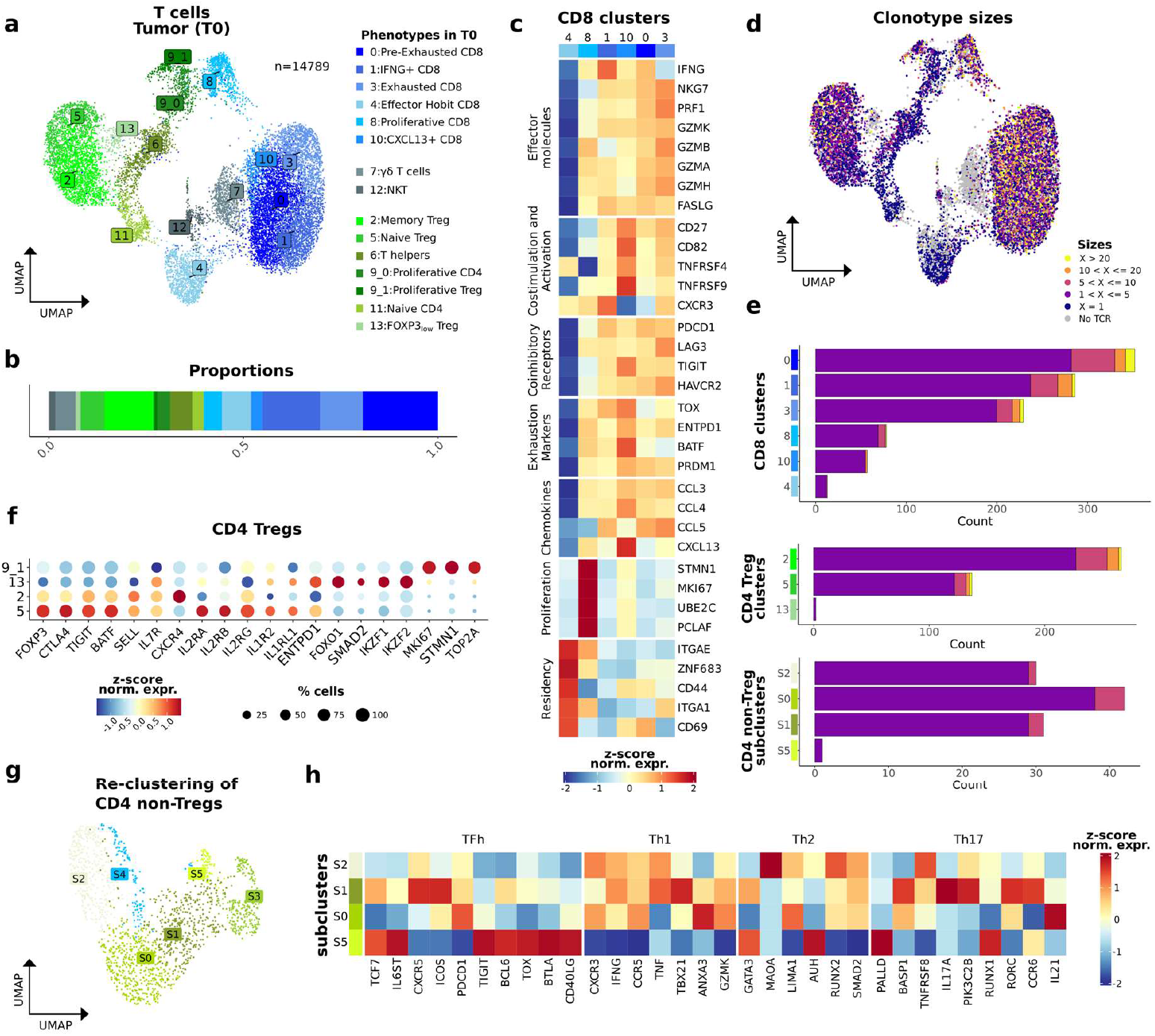
Profile of T cells in the primary tumor microenvironment (T0). (**a**) UMAP with T cell types and states, including CD8 (blue), CD4 (green) and non-conventional (gray) T cells. (**b**) Bar plot displaying the proportions of the T cell populations in the TME. (**c**) Heatmap of CD8 clusters, displaying pseudobulks of normalized aggregated counts for each cluster, transformed into z-scored values. The selected genes represent markers of functional T cell states. (**d**) UMAP of T cell (a), colored by the clonotype size of each T cell clone. (**e**) Bar plots presenting the total number of clonotypes, and indicating the clonotype size per cluster (divided into T cell types; i.e., CD8 (top), CD4 Tregs (middle) and CD4 non-Treg subclusters (bottom)). TCR singlets were omitted in this plot. (**f**) Dot plot of CD4 Treg clusters, displaying pseudobulks of normalized aggregated counts for each cluster (colors indicate z-scored values and dot size represents the percentage of cells expressing the gene). Selected genes represent marker genes of the respective clusters. (**g**) UMAP of clusters 9_0, 6, and 11 (a) after subclustering of non-Treg CD4 cells, identifying proliferative CD8 (S4), naive CD4 (S3) and four T helper subtypes (S0,S1,S2,S5). (**h**) Heatmap of the subclusters S0,S1,S2 and S5 (g), displaying canonical markers and annotation of T helper cell subtypes.

### Clonal tracking of anti-tumoral T cell activity in the blood

Tracking TCR clonotype dynamics during tumor progression or treatment response is notoriously challenging, since repeatedly sampling of the primary tumor is clinically complex due to patient frailty and limited access. Thus, we considered whether a deep characterisation of circulating T cells in the peripheral blood could, in addition to facilitating longitudinal tracking of tumor TCR clones, identify clonal TCR expansion upon ICI. Therefore, we employed deep single-cell T cell sequencing (OS-T) to profile 2,003,861 CD3+ T cells isolated from the patient’s blood before ICI treatment (T1), during chemotherapy and immunotherapy (T2-T5) and with immunotherapy alone (T6-T7; **Figure 1b**), covering a total of 12 months after the initial MRT diagnosis. From the 47,804 expanded T cell clones (>1 copy) in the TME, 4,594 (9,61%) were detected in the blood. Expanded TME-resident clones represented 0.4% of T cell clonotypes of the circulating immune repertoire (4,594/1,085,362), highlighting the need for deep T cell sequencing for clonal tracking. Intriguingly, TME-resident clones of distinct immuno-phenotypes were represented in the blood at different time points, reflecting different phases of the anti-tumor immune response (**Figure 3a**). While predominantly TME clonotypes of CD4 Treg cells were detected in the blood before ICI treatment (T1), these proportions strongly shifted towards the representation of cytotoxic CD8 cells after treatment initiation (T2-3), and pointing to the activation of a systemic anti-tumor response. Surprisingly, later time points during ICI treatment (T4-T7) revealed a shift to CD4 T helper cells, suggesting the formation of a stable adaptive immunity against the tumor.

**Figure 3:**
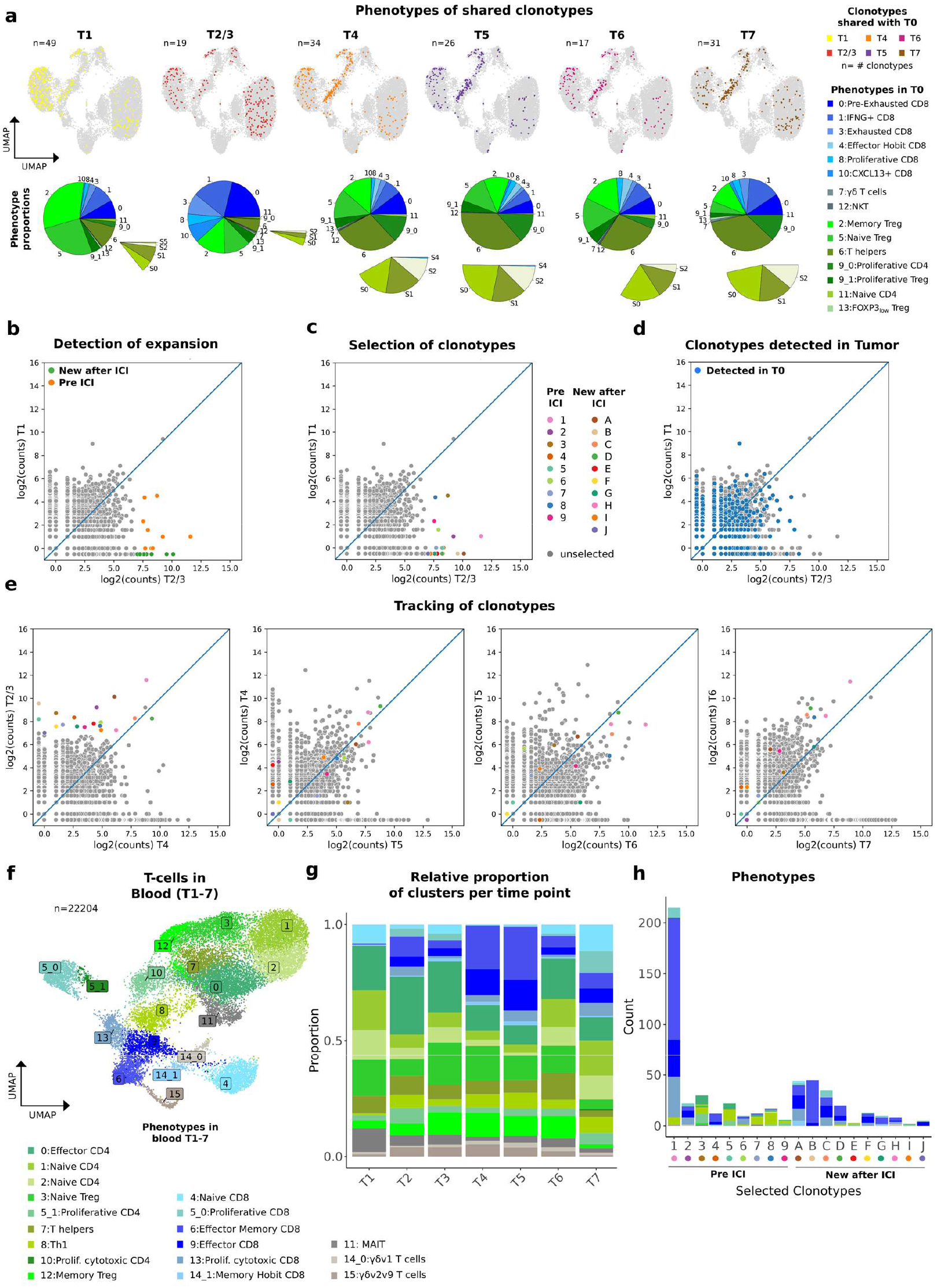
Spatio-temporal tracking of TME-resident and ICI-induced T cell clonotypes in the blood. (**a**) UMAP of TME-resident T cells (T0, Figure 2a) highlighting the TCR clones detected in the blood using deep single-cell T cell sequencing (OS-T). The overlap of the TME-resident clones and the blood is split by sampling time point (T1-7), illustrating the evolution to T cell representation in circulating T cells during the anti-tumor response of the patient. The pie charts illustrate the proportions of each T cell subpopulation from the TME that is represented in the blood over time. (**b**) Scatter plot showing clonotype counts of T cells in blood, comparing before (T1) and after (T2/3) ICI treatment initiation. The colored clonotypes represent significantly expanded T cells clones after ICI treatment, split in pre-existing (orange) and newly expanded (green) clones. (**c**) Scatter plot as in (b), highlighting T cell clones found in the TME (T0, blue). (**d**) Scatter plot as in (b), individually highlighting significantly expanded clonotypes that are used for further tracking and validation, split by pre-existing (1-9) and newly expanded clones (A-J). (**e**) Tracking of the ICI-induced clonotypes during consecutive time points (T2/3-T7). (**f**) Integrated UMAP of circulating T cells in the blood (all time points), clustered and annotated using gene markers. (**g**) Stacked bar plots of the proportional composition T cell population across time points. (**h**) Stacked bar plot of the total cell count per significantly expanded clonotype (b), colored by cell subpopulation (f).

Considering that ICI triggers anti-tumor immunity of both pre-existing and newly activated T cell clones in adult tumor patients, we then quantified T cell clonal expansion during the first cycles of ICI (T1/T2-3), using clonal activity dynamics as a proxy for ICI efficacy (**Figure 3b**). Due to the patient’s young age and the lack of pathogen exposure, as well as the prior chemotherapy, circulating T cells were mostly represented by unique clonotypes (n=1; 82.53%), with few clonotypes showing moderate expansion profiles (n>1; 17.47%, 2 median expansion count, **Extended Data Fig. 3**). This is in stark difference to adult immune repertoires profiled with the same technology, showing large, expanded clones representing past immune challenges (**Extended Data Fig. 3**). Nevertheless, we identified significantly expanded T cell clones after the first cycles of ICI, with the expansion of both pre-existing (clones 1-9) and newly activated T cell clonotypes (clones A-J), suggesting the combination of prior and novel clonotypes to be the driver behind the therapy success (Noise model, p<0.05; **Figure 3c**). Of note, 5 out of 19 of significantly expanded clones were also detected in the TME (clones 6,8,9,C,H; **Figure 3d**). Most importantly, 13 of 19 clones were still detected 12 months after therapy initiation (T7), indicating a long-lasting ICI-induced T cell memory in a context where persistence of anti-tumor T cell immunity is highly relevant (**Figure 3e**).

Detecting ICI-induced T cell activity that resulted in stable clonal memory prompted us to perform immuno-phenotyping of the respective clones. Single-cell transcriptome and TCR sequencing from the respective time points (T1-7) displayed unique features of the pediatric immune system (**Figure 3f**). Before ICI treatment, we found mainly naive CD4 and Treg population (**Figure 3g, Extended Data Fig. 4a** and **b, Supplementary Table 6**). After ICI treatment, T cell composition shifted to increased levels of cytotoxic and proliferating CD8 T cells, the former being conserved in subsequent time points. In addition to the cell composition, we observed an increase in expanded T cell clonotypes after ICI treatment (**Extended Data Fig. 4c**), together indicating the active mounting of a systemic adaptive immune response. The previously identified ICI-induced T cell clonotypes represented both CD4 and CD8 subtypes when considering only pre-existing clones (1-9), but mostly CD8 cells for newly activated clonotypes (A-J, **Figure 3h**). Here, clones were actively proliferating and showed clear signs of cytotoxicity and exhaustion. Together, we identified T cell clones expanding after ICI treatment that subsequently converted into a stable immune memory of circulating T cells. These clones represented active immuno-phenotypes, further suggesting their role in the CR of the patient and the long-lasting therapeutic effect.

**Figure 4:**
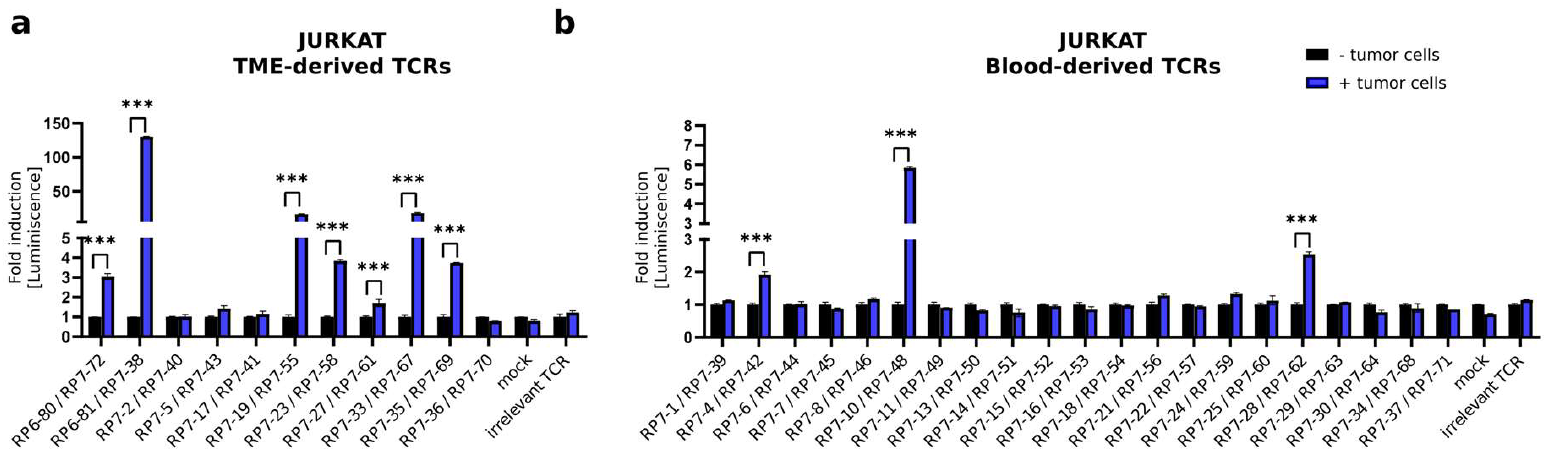
Validation of tumor reactivity of candidate TCRs in Jurkat cell model. α/β TCR-KO Jurkat cells transfected with tumor TME-derived **(a)**, or blood-derived **(b)** candidate TCRs were co-cultured with the patient’s tumor cells in a 1:1 ratio, and NFAT activation was measured using reporter luminescence signal. Mock-transfected Jurkat cells, or Jurkat cells transfected with a patient-irrelevant TCR, were used as negative controls. Data represented is mean luminescence signal +/- SEM, and is normalized to the “no tumor” condition for each tested TCR.

### TCR-engineered T cells for personalized cell therapy

Given the evidence for a strong anti-tumor immune response and the abundance of tumor-reactive T cells in the patient, we next sought to simulate a personalized T cell therapy for adoptive cell transfer (ACT) using TCR-engineered T cells as infusion product. Therefore, we used a well-established model system (Jurkat cells) and the patient’s own T cell pool from PBMCs (anti-CD3 and IL-2 expanded) to engineer T cell products using tumor-reactive TCR candidates. These candidates were derived through three complementary strategies to cover tumor-derived (TILs) and therapy-induced (blood) clonotypes. Tumor-derived clones were selected using 11 of the most expanded TILs (**Extended Data Fig. 5**). 21 therapy-induced clones were selected based on their expansion upon ICI treatment (both pre-existing and new clonotypes). Noteworthy, the deep single-cell TCR sequencing (OS-T) allowed the detection of both TCR chains (TRA/TRB) as an input for subsequent T cell engineering of candidate clonotypes. For rapid and efficient T cell engineering, we transiently transfected (codon-optimized) TCR alpha and beta chain mRNA. For this, the human constant TCR region was replaced by a mouse sequence to prevent the pairing with endogenous chains, as previously described^26^. The TCR-engineered T cells were then exposed to a tumor cell line derived from the patient’s primary MRT for activity testing.

Here, both tumor-derived and therapy-induced clonotypes showed reactivity against the tumor cells. Most strikingly, this strategy validated the anti-tumor activity of the majority of tested T cell clonotypes (7 out of 11) from the tumor in either the autologous and T cell lines model, representing CD8 and CD4 T cell clones (**Figure 4a, Extended Data Fig. 6a**). This is of utmost therapeutic importance considering the challenge to identify tumor-reactive clonotypes in adult tumors, where the TME contains high frequencies of bystander T cells reactive to viral antigens^27^. Our results point to the natural enrichment of tumor-reactive T cells in the TME, likely due to the naive characteristics of the early developing immune system. In turn, this highly facilitates the identification of TCR candidates for TCR-T therapy, but also allows us to speculate unselected TIL infusion products to be more efficient in the pediatric as compared to the adult context. Moreover, we also validated the tumor-reactivity of therapy-induced T cell clonotypes from the blood (3 out of 21), confirming T cells in circulation to serve as a resource to identify TCR-T candidates and highlighting deep single-cell TCR sequencing as a powerful tool for candidate selection and prioritization (**Figure 4b, Extended Data Fig. 6b**).

## Discussion

We identified and validated tumor-reactive TCR sequences from the TME and from T cells in circulation to serve as candidates for a personalized TCR-T therapy of an MRT patient. While the identification of TCRs from the TME for TCR-T treatment has been shown previously, and even entered Phase 1 clinical trials^28^, the selection of TCR candidates is cumbersome and highly prone to false-positive sequences. This is mostly due to the fact that in adult tumors the relative frequency of tumor-reactive clonotypes can be very low and clonal expansion and gene expression features are imperfect in distinguishing tumor-reactive from bystander T cells. In contrast, we were able to validate the tumor-reactivity of the majority of clones selected from the TME of this childhood cancer patient, likely due to the lack of bystanders in the immature immune system of this patient. If replicated across more childhood cancer cases, such accumulation of tumor-reactive T cells can have profound impact on the success of TIL products and the selection of TCR-T candidates, with the natural enrichment strongly increasing the frequency of tumor-killing T cells and true-positive TCRs, respectively. Likewise, the lack of validation of ICI-induced T cells in the blood to serve as a valuable resource for TCR selection for therapeutic use has geared the selection focus to the TME. However, the longitudinal tracking through liquid biopsies has tremendous potential, increasing sample accessibility and capturing T cell response dynamics over time. The latter is particularly important to extend profiling beyond the static snapshot obtained from the initial primary tumor resection, thus integrating data throughout tumor adaptation to therapy, as well as covering T cell activity at sites distant from the primary tumor. Consequently, identifying T cells from the blood can widen the application spectrum for TCR discovery and allows TCR-T design to be dynamically adapted together with the tumor evolution during treatment. Here again, the immature nature of the developing immune system provides an advantage when selecting TCR-T candidates. Specifically, the lowly expanded T cell memory in circulation allows the ICI-driven anti-tumor response to exceed clonotype sizes of pre-existing clones, facilitating the detection of tumor-reactive TCR candidates. It is also worth speculating that the candidate selection is supported by the combined treatment with chemotherapy, particularly cyclophosphamide, which contributes to the resetting of the adaptive immune repertoire and, hence, amplifies the signal of subsequent ICI-induced T cell activity. The validation of an unprecedented fraction of ICI-induced TCRs in this work strongly supports this notion. Moreover, we likely underestimate the potential of therapy-derived T cell clonotypes in the T cell engineering assays. These clonotypes were identified from peripheral blood samples obtained weeks after the resection of the primary tumor, from which the tumor cells for *in vitro* validation were derived. If they reacted to novel tumor antigens derived from metastatic sites or chemotherapy-induced antigen spreading, their activation upon exposure to the primary tumor was not expected. Hence, they might still be potent cell therapy candidates for a relapsed tumor, matching antigens exposed during tumor evolution, a specificity that could be validated by challenging respective TCR-products with biopsy/resection material at relapse.

TCR-T therapy represents a powerful extension of the cell therapy portfolio, for which CAR-T therapy is the most prominent example^29^. TCR engineered cells have the advantage by also targeting intratumoral epitopes through their display as peptides in the context of MHC complexes on the cell’s surface. TCR-Ts can target common cancer epitopes, such as the recently FDA approved therapy using engineered T cells against MAGEA4^30^ for the treatment of advanced synovial sarcoma. In addition to tumor-associated antigens, recurrent mutations can be targeted, as shown for hotspot *KRAS* G12D mutations in pancreatic cancer^18^. Given the recent successes and approvals in TCR-T therapies and the general advances in cell therapy development and manufacturing, a personalized strategy is within reach. Here, we propose a framework that overcomes previous hurdles in the selection of TCR candidates. Achieving high TCR validation frequencies, we propose TCR selection from both the TME and the blood to be viable strategies covering different clinical scenarios to identify and track clones through needle and liquid biopsies, respectively. A personalized infusion product can also be envisioned as a multi-TCR product, similar to the pooling strategy applied for personalized cancer vaccines^31,32^. Pooling multiple TCRs might offer a more robust and durable response against the tumor through the targeting of different antigens and by representing diverse receptor-epitope affinities with different activation and exhaustion profiles.

The here tested cell engineering strategy, through the application of transient mRNA transfection, has additional benefits as compared to the standard practice of stable retro- and lentiviral cell transduction. Transient strategies using mRNAs would strongly reduce the risk for long-term off-target effects, which could lead to lifetime risk of comorbidities, especially important in the context of treating children with cancer. In addition, considering recent advances of *in vivo* cell engineering strategies for viral epitope (Moderna) or targeted CAR-T expression^33^, an adaptable, personalized TCR-therapy strategy can be envisioned. Besides practical and financial advantages compared to *ex vivo* approaches, *in vivo* cell manipulation would also allow dose-escalation and repeated boosting. Such *in vivo* therapies, for which mRNA encoding for TCR sequences is encapsulated into T cell-targeting nanoparticles, have already been successfully tested in preclinical models^34^.

A key part of this case is the high presence of CD8+ T cells infiltrating the tumor, indicating an active anti-tumor immune response. This clear immunogenic tumor profile highlights the potential for the broader application of ICI in rare childhood cancers and for next-generation therapies (e.g. autologous T cell therapies). A possible cause for the high immunogenicity, despite the generally low mutation burden in MRTs, is the expression of endogenous viruses. This idea aligns with previous studies suggesting that the loss of SMARCB1 in MRTs may lead to the derepression of endogenous retroviruses, which trigger an immune response^12^. In fact, a viral-like immune response may be stronger than a neoantigen-driven anti-tumor response, which could help to explain the complete remission of the patient.

In summary, we validate deep single-cell TCR profiling of both TME and peripheral blood as a valuable approach for identifying tumor-reactive T cells in a low-TMB childhood cancer patient and establish a framework and proof-of-concept for TCR engineering for personalized TCR-T therapy.

## Methods

### Ethical approval

Patient material was included in the Sant Joan de Déu Barcelona Children’s Hospital Biobank (BHISJDI) for Research after written informed consent from the patient’s legal guardians. Access to said samples granted by the Hospital Sant Joan de Déu Drug Research Ethics Committee (CEIm) under approval number PIC-30-21. Biological material from the patient safeguarded in the Biobank will be governed according to the Biomedical Research Law (Law 14/2007, of July 3, 2007), and by the provisions of Royal Decree 1716/2011, of November 18, 2011.

### Sample collection

Fresh tumor surgical resection material was gently minced into small fragments (0.5 - 2 mm) using sterile scalpels. The resulting tumor pieces were cryopreserved in Synth-a-Freeze Cryopreservation medium (Gibco) following the manufacturer’s instructions. A small fraction of freshly processed tumor pieces were used to establish patient-derived tumor cells (see below).

Peripheral blood mononuclear cells (PBMCs) were isolated directly from fresh blood by density gradient centrifugation using Lymphoprep (Stemcell, Germany) in accordance with the manufacturer’s instructions. Subsequently, the PBMCs were cryopreserved in a freezing medium composed of 50 % RPMI 1640 supplemented with fetal bovine serum (FBS), 40% FBS, and 10 % DMSO. The cryopreserved PBMCs were stored at −80 ºC in a Mr. Frosty for 24 hours before being transferred to liquid nitrogen storage.

### Histopathological analysis

Histopathological evaluation of the surgically removed specimen was performed by trained pathologists from the Anatomical Pathology Unit at the Sant Joan de Déu Barcelona Children’s Hospital to confirm diagnosis, as well as specificity and localization of IHC staining. Surgically resected tumor tissue was fixed in 10 % neutral buffered formalin, paraffin-embedded and cut into 3 μm sections. IHC staining was performed in either a Leica Bond-Prime (Leica, USA) or Dako Autostainer Link 48 (Dako, USA) systems. The primary antibodies used were CD3 (IR50361, Agilent), INI1 (612110, BD; Bond-Prime), and PD-L1 (M365329, Dako). Stained immunohistochemistry sections of the primary tumor lesion were scanned using a high-resolution NanoZoomer 2.0 HT (Hamamatsu).

### Establishment of patient-derived tumor cells

Patient-derived tumor cells were established from the fresh, surgically resected primary tumor. Briefly, fresh tumor biopsy was mechanically dissociated using sterile scalpels, followed by enzymatic digestion with Trypsin-EDTA (0.25 %; Gibco) at 37 ºC for 5 minutes under agitation. After trypsin inactivation, the mixture was filtered through a 100 μm cell strainer and centrifuged at 300 x g for 5 minutes. Cells were cultured in tumor stem media (TSM) consisting of 1:1 mixture of DMEM/F-12 (Gibco) and Neurobasal-A medium (Gibco), 10 mM HEPES solution (Gibco), 1 mM Sodium Pyruvate (Gibco), 1x MEM Non-Essential Amino Acids Solution (Gibco), 1x GlutaMAX supplement (Gibco), 1x Antibiotic-Antimycotic (Gibco) supplemented with 10 % heat-inactivated fetal bovine serum. Cells were maintained in a humidified incubator at 37 ºC, 3 % O2, 5 % CO2, and media was changed every 48 h. Cells were routinely tested for mycoplasma contamination using standard PCR. Tumor cells used for the evaluation of tumor-reactive TCR candidates were maintained below passage 5 to preserve subclonal heterogeneity.

### Primary T cell isolation

PBMCs isolated from the patient (1–1.5 × 10^6^ cells/mL) were cultured in RPMI medium with Dynabeads® (ThermoFisher) at a 1:1 bead-to-cell ratio and supplemented with 60 U/mL recombinant IL-2. Cultures were incubated at 37 °C in a humidified CO^2^ incubator. Cells were examined daily for morphological changes and clump formation. Cell counts were performed twice a week, with cultures split to a density of 0.5–1 × 10^6^ cells/mL when the cell density exceeded 2.5 × 10^6^ cells/mL.

### TCR cloning and tumor reactivity validation

Protocol followed in here was performed as previously described by the authors (Pétremand 2024). Briefly, Antitumor reactivity of TCRs was assessed by transferring RNA encoding TCRαβ pairs into recipient activated T cells and a Jurkat cell line (TCR/CD3 Jurkat-luc cells (NFAT), Promega) stably transduced with human CD8α/β and with TCRα/β knocked out via CRISPR. Electroporated cells were co-cultured with interferon-γ (IFNγ)-treated autologous tumor cells. Tumor reactivity was evaluated by measuring CD137 upregulation in T cells and bioluminescence in Jurkat cells.

### Whole-genome sequencing

Whole genome sequencing (WGS) data was generated for tumor tissue and blood. Briefly, the tumor and blood samples were sequenced at 120X and 30X coverage respectively (150bp paired-end). Raw reads were processed with nf-core/sarek workflow (v3.4.4)^35^ using strelka2, mutect2 and freebayes variant callers with default parameters. Tumor mutation burden was calculated considering high-confidence mutations present in at least 2 of the 3 variant calls. Coding mutations were subset using annotated exons from protein coding genes (gencode v45).

### Sample preparation for single cell profiling

PBMCs samples were thawed, centrifuged at 350 x g for 5 min at 4 ºC and pellets resuspended in appropriate volume of PBS (Thermo Fisher Scientific) supplemented with 0.05% BSA (Miltenyi Biotec). Samples were filtered with a 40 um cell strainer (PluriSelect) and stained with trypan blue to assess final cell suspension concentration and viability, using a TC20™ Automated Cell Counter (Bio Rad). Cryopreserved tumor tissue was rapidly thawed at 37 ºC and subsequently processed to obtain a single-cell suspension. Briefly, tumor chunks were equilibrated to RT for 10 minutes in pre-warmed RPMI 1640 medium (Gibco) supplemented with 10 % FBS, 1 % Pen/Strep and 1x GlutaMAX supplement. Tumor chunks were washed twice with Hank’s Balanced Salt Solution (HBSS; Gibco) and mechanically dissociated before enzymatic digestion in 1.25 mg/mL Collagenase A (Sigma), 50 ug/mL DNAse I (Sigma) in HBSS at 37 ºC for 20 minutes, with regular gentle agitation to facilitate disaggregation. Cells were then washed twice in ice-cold PBS and passed through a 100 μm cell strainer to obtain a single-cell suspension for subsequent FACS-sorting.

Single-cell suspension obtained from the tumor tissue were stained with anti-CD45 (PerCP/Cyanine5.5 anti-human CD45 Antibody, Biolegend) and anti-CD3 (PE/Cyanine7 anti-human CD3 Antibody, Biolegend) antibodies for 20 minutes at 4 ºC protected from light. FACS sorting was performed using the Melody FACS flow cytometer (Becton Dickinson, Franklin Lakes, NJ) to separate the immune compartment from the TME, and to positively select the lymphoid T cell fraction. After sorting, CD45-positive and CD3-positive cells were centrifuged at 400 x g for 7 minutes at 4 ºC and counted with a TC20™ Automated Cell Counter (Bio-Rad Laboratories, S.A), before proceeding to single cell RNA sequencing.

### Library preparation and single-cell RNA-sequencing

Samples were loaded onto the Chromium X instrument (10X Genomics) and encapsulated for a target cell recovery of 20,000 cells, using the Next GEM Single Cell 5’ HT Reagent Kits v2 (10X Genomics, PN-1000356). All samples were loaded in duplicate except time point 0 (FACS sorted cell fractions from the tumor) and time point 5 (PBMCs). Sequencing libraries were prepared according to the manufacturer’s instructions. Briefly, after GEM-RT cleanup, cDNA was amplified over 13 cycles, purified, and quantified on an Agilent Bioanalyzer High Sensitivity chip (Agilent Technologies). Human T cell receptor (TCR) sequences were enriched from the amplified full-length cDNA using the Chromium Single Cell Human TCR Amplification Kit (PN-1000252). To construct the gene expression (GEX) and TCR libraries, 30 to 150 ng of cDNA were fragmented, end-repaired, A-tailed, and sample-indexed using the Chromium Single Cell 5’ Library Construction Kit (10X Genomics, PN-1000190) and the Dual Index Kit TT Set A (10X Genomics, PN-1000215). Size distribution and concentration of the libraries were verified on an Agilent Bioanalyzer High Sensitivity chip and finally sequenced on a NovaSeq 6000 system (Illumina), aiming for approximately 40,000 and 10,000 read pairs per cell for the GEX and TCR libraries, respectively. The sequencing conditions were 28 bp (Read 1) + 10 bp (i7 index) + 10 bp (i5 index) + 90 bp (Read 2).

### OS-T (Single-cell TCR sequencing)

For OS-T assay, cryopreserved PBMC were thawed by immediate immersion of the vials in a 37ºC water bath, and then washed in complete media. After washing PBMC in PBS with 0.05% BSA, samples were run with the Omniscope (Omniscope Inc., Barcelona, Spain) proprietary single-cell technology OS-T to generate sequencing libraries.

### OS-TCR (Single-molecule TCR sequencing)

Four snap-frozen tumor samples, each consisting of 5-6 sections with a thickness of 50 micrometers, were employed for RNA isolation. RNA isolation was performed using the RNeasy Plus Mini Kit (QIAGEN). The isolated RNA served as the input material to perform the Omniscope (Omniscope Inc., Barcelona, Spain) single-molecule OS-TCR assay to generate sequencing libraries.

## Data processing

### 10X Genomics data (TME and PBMCs time point samples)

Raw gene expression (GEX) and V(D)J repertoire data (FastQ files) were processed using the CellRanger (v7.0.0) multi pipeline [https://www.10xgenomics.com/support/software/cell-ranger/latest/analysis/running-pipelines/cr-5p-multi] and mapped to the human GRCh38 reference genome (GENCODE v32/Ensembl 98). All the data analysis generated with 10x Genomics has been analyzed within the Seurat (v4.0.5) framework in R (v4.1.1). Detailed list of the packages used on each step of the analysis are reported at the end of the notebooks containing the code that are provided in the project’s Github repository.

### Quality Control (QC)

We uploaded the *sample_filtered_feature_bc_matrix* yield by CellRanger of each library into a Seurat object (see Supplementary Table 1). First, for each library matrix, we filtered out genes with 0 counts. Secondly, to filter out low-quality cells, we applied different thresholds to the GEX data for each library based on the distribution of total UMIs and unique gene counts. Specifically, we excluded cells with fewer than 1,000 UMIs and capped the maximum UMI threshold according to each library’s distribution, setting it at the 0.99 quantile, which ranged between 14,000 and 37,000 UMIs. Similarly, we removed cells with low unique gene counts, setting a minimum threshold at the 0.10 quantile and a maximum at the 0.99 quantile of the unique gene distribution per library. At this point, we used permissive thresholds since the clustering and marker analysis downstream (see upcoming sections) further provided more evidence to remove low quality cells. As part of QC, we ran Scrublet (v0.1.0) and added the doublet detection score yield by this tool to the metadata of our Seurat objects per library.

### Pre-processing and initial analysis

At this point, data of libraries belonging to technical replicates from the PBMCs timepoints were merged into a single Seurat object. We normalized cell counts by dividing by the total gene counts for each cell, scaling by a factor of 10,000, and then applying a log transformation. Feature selection was performed modulating the mean and the variance of the log-normalized values using *modelGeneVar* function from scran (v1.22.1) and setting an FDR threshold of 0.05 at *getTopHVGs* function to obtain a list of highly variable genes (HVGs). T cell receptor genes were excluded from the list of highly variable genes (HVGs) to prevent them from influencing downstream clustering, as they can cause certain similar cells to cluster a part. HVGs lists obtained were added as variable features in the corresponding Seurat objects and used for scaling the data (z-score transformation) and for computing Principal Component Analysis (PCA). Principal components (PC) were selected for each Seurat object based on the Elbow Plot of the standard deviation (see HTML reports on repository for each case). Uniform Manifold Approximation and Projection (UMAP) was used to visualize data. Lastly, we computed the cell cycle score using the gene list provided by the function *cc*.*genes* and added cycle ‘phases’ (G1, G2M, and S) to the metadata. At this stage, we observed that for certain PBMC time-point samples, hemoglobin genes associated with erythrocytes appeared among the top highly variable genes (HVGs) and were highly expressed in some cells, as visualized with the *FeaturePlot* function. In these cases, we retained only cells with zero counts for the genes *HBM, HEMGN*, and *HBG2*, then pre-processed these subsets following the same steps as described above.

We applied Louvain clustering by first building a k-nearest neighbors (KNN) graph and then defining clusters at varying resolutions using Seurat’s *FindClusters* function. For each dataset, we selected the resolution that best represented the major cell types. To identify these major cell types, we performed a “one-vs-all” differential expression analysis to find markers specific to each cluster, using Seurat’s *FindAllMarkers* function with the Wilcoxon rank-sum test (*test*.*use = “wilcox”*).

In this initial clustering, we identified additional low-quality cells, erythrocytes, and doublets. Doublets were characterized as clusters containing mixed cell type markers and showing high Scrublet scores. After removing these clusters, we re-processed the subsets using the same steps as previously described, followed by a final clustering. We then manually annotated the clusters by examining the expression levels of canonical gene markers for major cell types.

## Data analysis

### Immune cells in Tumor sample (T0)

After processing the single-cell GEX data of the sorted cell fractions (CD45+ and the CD3+ enriched) of the tumor sample (T0), we integrated these two to provide a general overview of the TME populations recovered in this study. We first merged the Seurat objects corresponding to these two sorted fractions and then normalized again as previously described. For variable features in the merged TME object, we used the union of HVGs identified in each fraction. Then, we scaled the data and computed PCA as described above. Based on the Elbow Plot of the standard deviation of the components, we selected the first 25 PCs for downstream analysis. To ensure robust integration of T cells across fractions, we applied Harmony (v1.0.3) to correct potential batch effects within the 25 PCs of the combined dataset (**Extended Data Fig.1a)**. Again, we used the Louvain algorithm for clustering based on a KNN graph and defined clusters with 0.2 resolution. By manually inspecting canonical marker genes for T cell populations, we used Seurat’s *FindSubCluster* function to achieve greater granularity within cluster 5 (**Extended Data Fig.1c**). We then run Seurat’s function *FindAllMarkers* with the default Wilcoxon rank sum test to obtain differential expressed genes between all clusters. Looking at the list of marker genes per cluster and the annotations previously defined during the pre-processing step (see above), we annotated them providing general cell types (see **Figure 1d, Extended Data Fig.1d**).

### T cells in Tumor sample (T0)

To achieve greater granularity in identifying T cells within the TME, we subsetted the CD45+ fraction specifically to T cells, based on the subtypes defined during the initial pre-processing. We joined this subset with the CD3+ fraction into a single Seurat object of all T cells of the TME. We computed the union between the HVGs list of the enriched CD3+ fraction and the ones obtained for the T cell subset of the CD45+ fraction again modulating mean and variance as previously described. We performed the same steps of scaling and computing PCA. Elbow Plot suggested considering the first 15 PCs this time. We harmonized the data of the fractions (using Harmony v1.0.3) correcting by fraction and found clusters as described before using a resolution of 0.8. Once more, we used Seurat’s *FindSubCluster* option to obtain more granularity on the proliferating cells of cluster 9 which helped us separate different cell types proliferating. Wilcoxon rank-sum test implemented in Seurat’s *FindAllMarkers* function provided a list of markers per cluster that we used to annotate the T cells (see **Figure 2a, Supplementary Table 4**). In order to further explore clusters 9_1, 6 and 11, we subsetted them and processed them together following the same steps as mentioned before. The first step involved selecting highly variable genes (HVGs) by combining the HVG lists obtained from the CD3+ fraction and the T cell subset of the CD45+ fraction. These HVGs were selected by modulating the mean and variance of the gene expression values as previously described. After selecting the HVGs, we log-normalized the data, scaled it, and performed PCA, using the first 10 PCs as indicated by the Elbow Plot. We computed the KNN graph and clustered with Louvain obtaining the 5 subclusters shown in **Figure 2g**. We computed marker genes per cluster and found that subcluster 4 (S4) are actually CD8 proliferative cells that initially clustered within 9_0 and that subcluster 3 (S3) are Naive cells (see **Supplementary Table 5**). The rest of the subclusters are different T Helper groups (see **Figure 2h**).

### T cells from PBMCs (T1-7)

After the initial pre-processing and clustering steps described earlier, we had obtained a separate Seurat object for each time point. We subset those to the clusters that we previously annotated as T cells during the pre-processing step. We then combined all blood PBMC samples into a single Seurat object and normalized cell counts by dividing by the total gene counts for each cell, scaling by a factor of 10,000, and then applying a log transformation. To define highly variable genes (HVGs) for this merged dataset, we selected genes that were identified as “variable” in at least 2 of the time points samples. We selected the first 25 PCs and computed a UMAP to visualize the merging of the time points. We decided to proceed with batch correction of those using Harmony and correcting by *library* as those were sequenced separately (see **Supplementary Table 1**). We computed the KNN graph and clustered by applying Louvain with resolution of 0.8. This initial clustering served to further detect low quality cells, doublets and a cluster with some hemoglobin genes. We removed those and followed the same steps as before, but this time selecting 20 PCs and using a resolution of 1 from where we obtained 15 clusters for annotation. Additionally, we used Seurat’s *FindSubCluster* function to further proliferate cells in cluster 5 and to gain more resolution in cluster 14. We computed the markers per cluster as previously described in the above sections and manually annotated the cells as shown in **Figure 3f**.

### Marker genes

To define CD8 cluster functions, we used the genes and the function names reported by Li et al., 2023^36^. We also added to the heatmap (**Figure 2c, Extended Data Fig.4a**), a curated list of genes representative of T cell tissue residency (Molodtsov and Turk 2018). CD4 cluster dot plots of **Figure 2f** and **Extended Data Fig.4b** contain genes obtained as top marker genes per cluster as previously explained but manually selected those that better help visualize the different cell types. In all the mentioned figures representing marker genes, we aggregate the raw counts per cluster and compute the logCPMs to normalize. For visualization purposes, we transform those values in z-scores.

### TCR repertoire analysis (10x Genomics)

From each library, we uploaded the *filtered_contig_annotations* file obtained with CellRanger into a list of contigs that we combined using scRepertoire (v1.12.0) setting the *filterMulti=TRUE* option of *combineTCR* function. This option selected the top 2 expressed chains in cell barcodes with multiple chains. In downstream analysis, we considered as a clonotype each unique beta chain nucleotide sequence. We added the combined contig information to the metadata of the corresponding Seurat object containing the T cells of either the TME or the merged PBMCs object.

### TCR repertoire analysis (Omniscope)

For quantitative TCR repertoire comparison and the selection of expanded T cell clonotypes after immunotherapy, TCR contig files (VDJ nucleotide sequences) were used as inputs for the comparative analysis between timepoints and for clonotype tracking. Identical contig IDs (V gene + J gene + CDR3 nucleotide sequence) for single cells (OS-T) or single molecules (OS-TCR) were grouped and quantified to determine the clonotype expansion for each timepoint. Log2 transformation of the counts was performed for visualization, since the clonotype counts follow a distribution with few highly abundant and many lowly abundant clones. Seaborn scatterplot python package was used to generate the scatter plots for the visualization of the repertoire counts between two timepoints. To select differentially expanded clonotypes between the timepoints before (T1) and after immunotherapy (T2/3) in the blood (OS-T), a proprietary noise model was applied (Omniscope Inc., Spain). The noise model generates a null distribution for each clonotype with a parameters to modify stringency.

## Declarations

### Data and code availability

The raw data (FASTQ files) for single-cell 10x Genomics data will be deposited in Gene Expression Omnibus (GEO) upon publication. Intermediate files such as count matrices of the GEX and TCR will be uploaded in GEO upon publication and accession codes will be provided. The code to reproduce the full analysis is hosted in a private Github repository and we will make it available upon publication.

### Author contribution

HH and AA conceived and supervised the project. IS and JLM performed all the computational analysis. IS analyzed all 10X Genomics data generated in this study. GC generated 10X Genomics datasets. JLM analyzed all Omniscope Inc. generated data. MG, MS and AMS generated Omniscope Inc. data. AA, HH, IS, JLM, AC and JCN interpreted the results. IS and JCN annotated the cells and provided phenotypes. AC and AUB established the patient-derived tumor cell line. JCN performed T cell isolation. AM, CS and AH, performed the T cell engineering experiment for validation of reactivity. CJRH, JPMP and JM provided the samples, the clinical information and are in charge of the ethical approvals. QR, DM, APR participated in providing additional insights and discussion. AA, HH and IC wrote the manuscript.

## Acknowledgements

IRB Barcelona is a recipient of a Severo Ochoa Centre of Excellence Award from the Spanish Ministry of Economy and Competitiveness (MINECO; Government of Spain) and an Excellence Institutional grant by the Asociacion Española contra el Cancer, and is supported by CERCA (Generalitat de Catalunya). We greatly appreciate the Market Solidari initiative for their continued support of our research endeavors. We are grateful to the Sant Joan de Déu Barcelona Children’s Hospital Biobank for Research and the Anatomical Pathology Department for their services. The authors would like to expressly thank the family of the patient who generously accepted the extraction and analysis of the samples. This study would not have been possible without this contribution.

