## Supplementary figures. for "Spatio-temporal T cell tracking for personalized TCR-T designs in childhood cancer"

**of**

### Extended Data Figure Legends (1-6)

**Extended Data Figure 1. TME clustering and annotation. (a)** UMAP in Figure 1d showing the integration of sorted fractions: the CD45+ and the enriched for CD3+ cells. (**b**) UMAP in Figure 1d with normalized expression of CD3D gene highlighting the T cells. (**c**) Resolution chosen for Louvain clustering was 0.2 with further sublclustering of cluster 5. (**d**) Dot plot showing canonical marker genes of each population for the annotation of clusters. Grey gradient of color means the average expression of the normalized expression per cluster and the size of the dot represents the percentage of cells expressing that gene.

**Extended Data Figure 2. CD4 T cell compartment of Tumor (T0).** (**a**) UMAP in Figure 2a showing on the left the log-normalized expression of FOXP3 on the right the CD40LG gene. (**b**) Violin plots showing the distribution of the log-normalized expression of FOXP3 (left) and CD40LG (right) of the clustering shown in Figure 2a.

**Extended Data Figure 3. Repertoire comparison in adult individual.** (**a**) Scatter plot showing clonotypes detected in blood sample (Timepoint 1) respect to a posterior blood sample (Timepoint 2), with the axes representing the total count of each clonotype on a log2 scale.

**Extended Data Figure 4. Single-cell profiles in blood time point samples.** (**a**) Heatmap of CD8 clusters displaying pseudobulks of normalized aggregated counts for each cluster, with z-scored values. The Y-axis represents a manually selected set of genes, grouped according to the functions they represent. (**b**) Dot plot of CD4 cell clusters showing pseudobulks of normalized aggregated counts for each cluster, with color indicating z-scored values and dot size representing the percentage of cells in the cluster expressing the gene. Genes are markers of each cell group. (**c**) Stacked bar plot showing the proportion of clonotypes at each time point, with color representing the size of each clonotype.

**Extended Data Figure 5**. **Tumor-reactive TCR candidates from expanded TILs.** (**a**) Same UMAP as in Figure 2a colored by the 11 expanded clonotypes selected for TCR-engineering from the Tumor sample. (**b**) Stacked bar plot showing the total counts (from single-cell data) and the phenotypes (color) of the 11 expanded clonotypes that were selected.

**Extended Data Figure 6**. **Validation of tumor reactivity of candidate TCRs in autologous PBMCs.** Patient-derived PBMCs transfected with tumor TME-derived (**a**), or blood-derived (**b**) candidate TCRs were co-cultured with the patient´s tumor cells in a 1:1 ratio, and 4-1BB expression (measuring activation) was quantified by flow cytometry. Mock-transfected PBMCs, or PBMCs transfected with a patient-irrelevant TCR, were used as negative controls. Data represented is mean 4-1BB-positive signal +/- SEM, and is normalized to the “no tumor” condition for each tested TCR. ND - not detected.

**Supplementary Table Legends (1-7)**

**Supplementary Table 1**. **Sample sequencing information.** This table contains the library information and the batches in which the 10x Genomics data has been sequenced.

**Supplementary Table 2**. **Canonical markers for annotation.** This table contains lists of genes used to annotate major cell types. These gene list are elaborated with in-house knowledge.

**Supplementary Table 3. Marker genes of Immune Cells at T0 (Tumor).** Table with the marker genes per cluster of the Immune cells of the Tumor. Clusters are those referred in Figure 1D and Extended Data Figure 1. Pval adjusted <0.05 and logFC>0.1.

**Supplementary Table 4. Table 4. Marker genes of T cells at T0 (Tumor).** Table with the marker genes per cluster of the T cells of the Tumor. Clusters are those referred in Figure 2a. Pval adjusted <0.05 and logFC>0.1.

**Supplementary Table 5. Marker genes of non-Treg CD4.** Table with the marker genes of the sublcustering of clusters 9_0, 6 and 11 referred in Figure 2g. Pval adjusted <0.05 and logFC>0.1.

**Supplementary Table 6. Marker genes of PBMCs T cells (T1-7).** Table with the marker genes of the T cells obtained from the PBMCs time point blood samples. Clusters refer to Figure3f . Pval adjusted <0.05 and logFC>0.1.

**Supplementary Table 7. TCR information of the candidates.** Table with the alpha and beta information of the TCRs used for the TCR engineering experiment referred in Figure 4. It also contains the correspondence with the labels used in other figures and the codes used in the validation experiment.

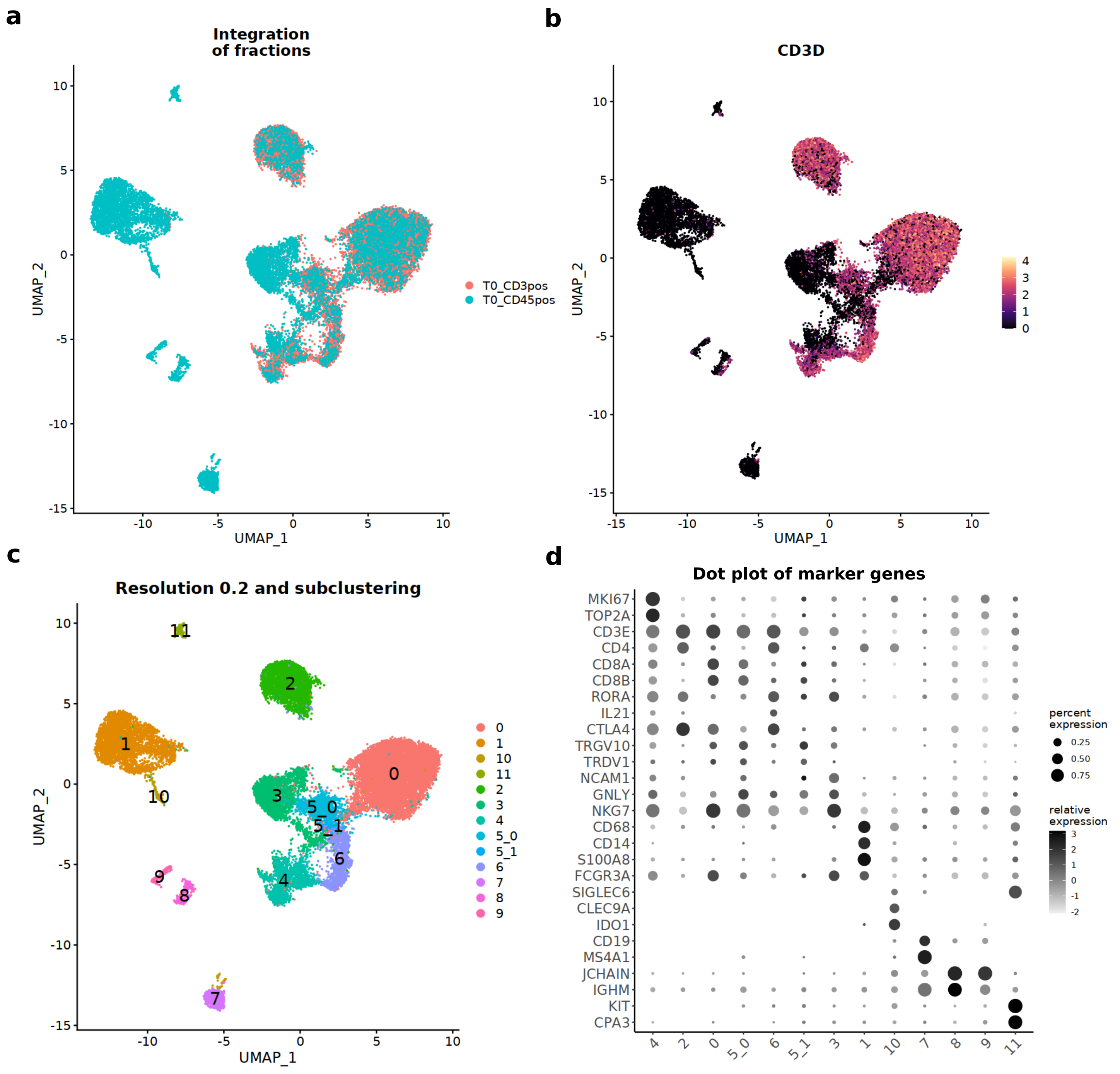


**Extended Data Figure 1.**

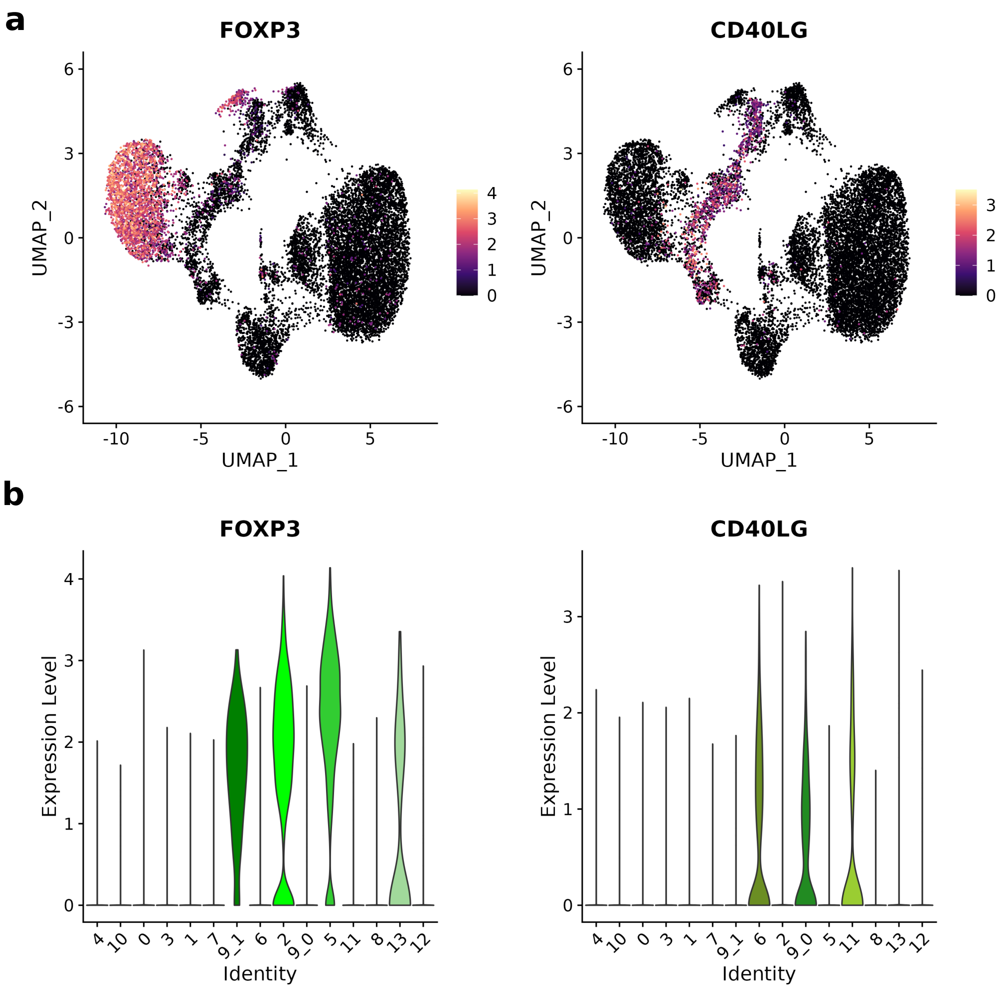


**Extended Data Figure 2.**

**a**

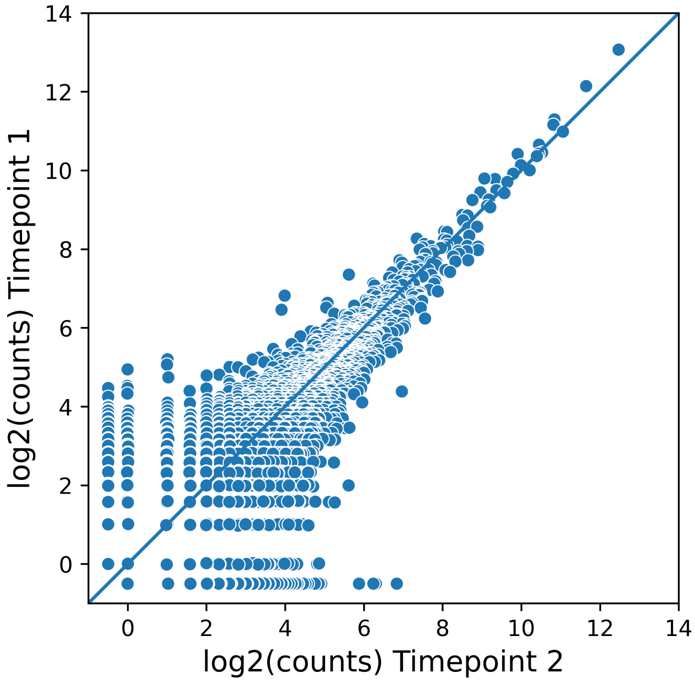


**Extended Data Figure 3.**

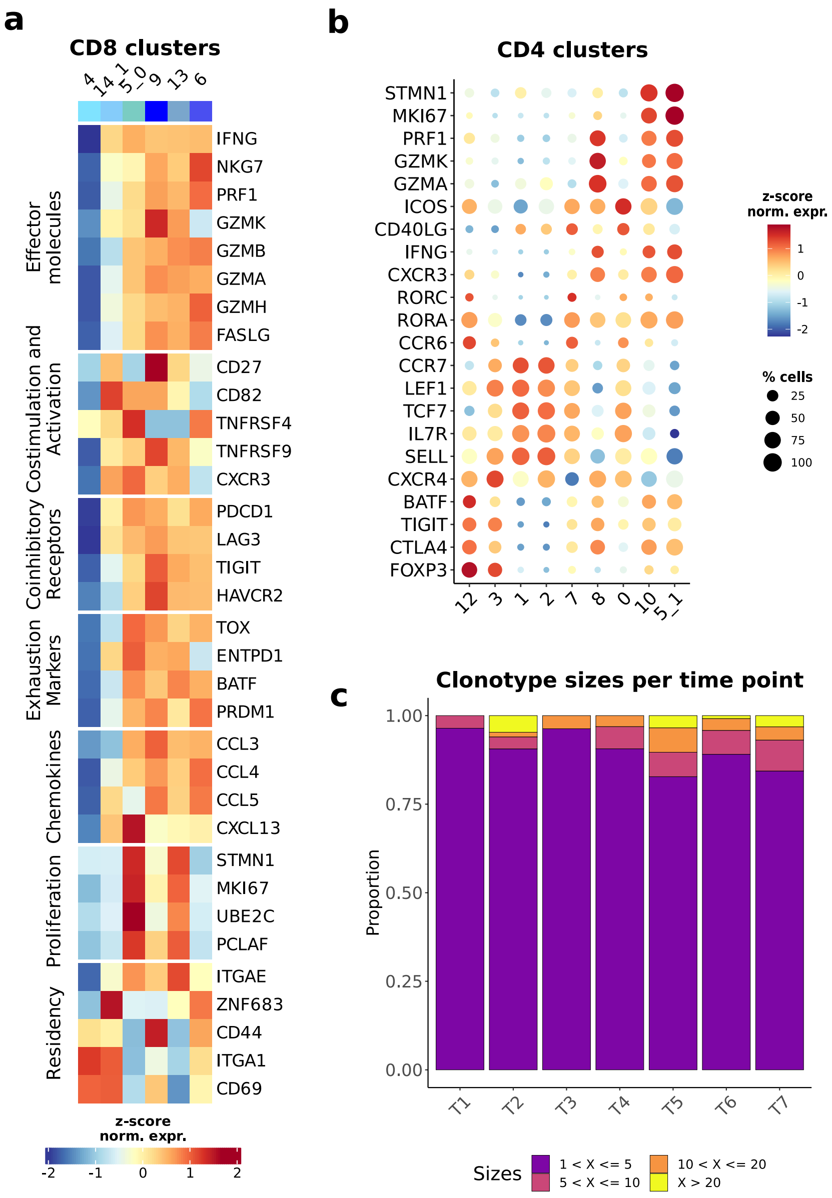


**Extended Data Figure 4.**

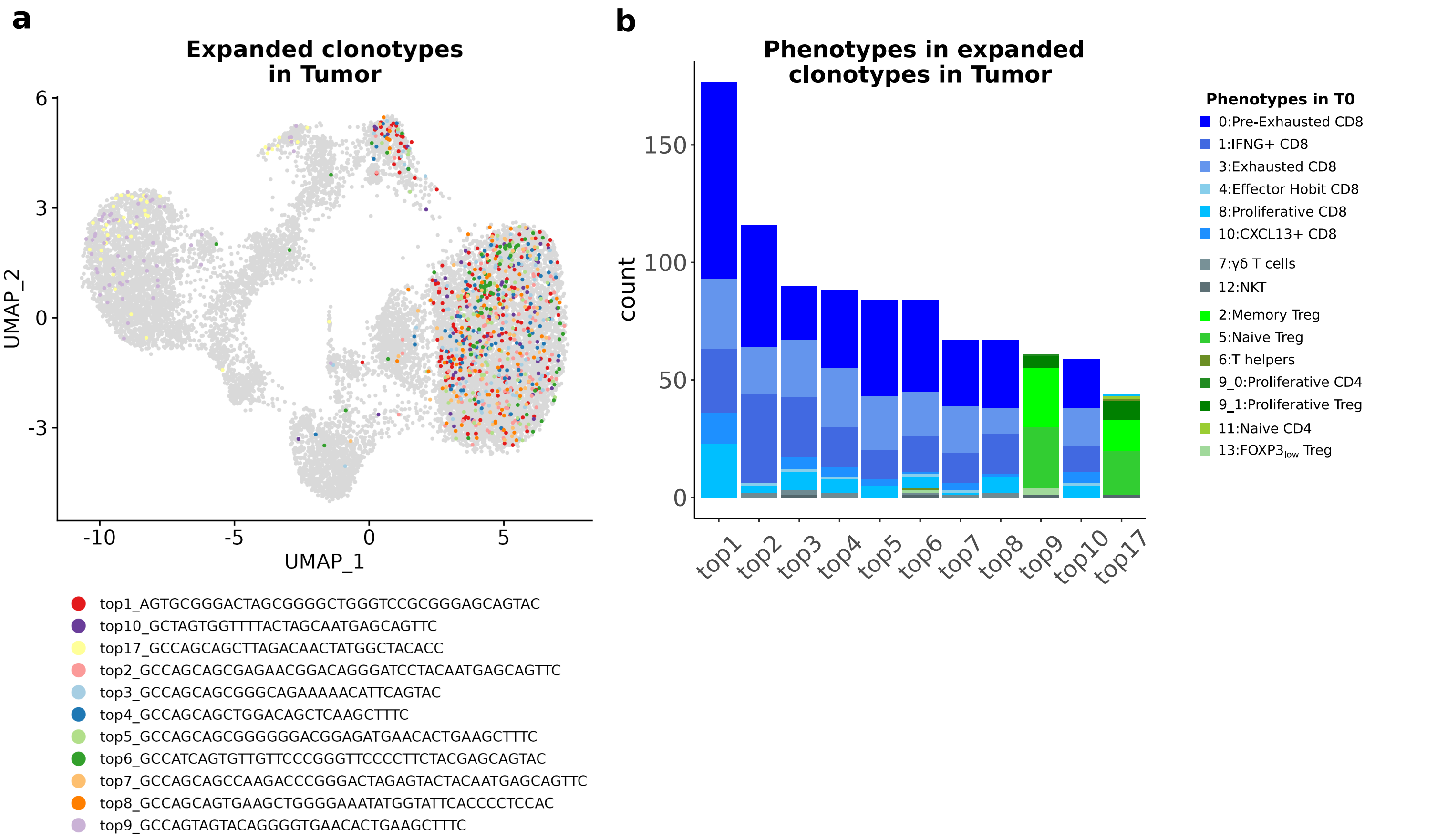


**Extended Data Figure 5.**

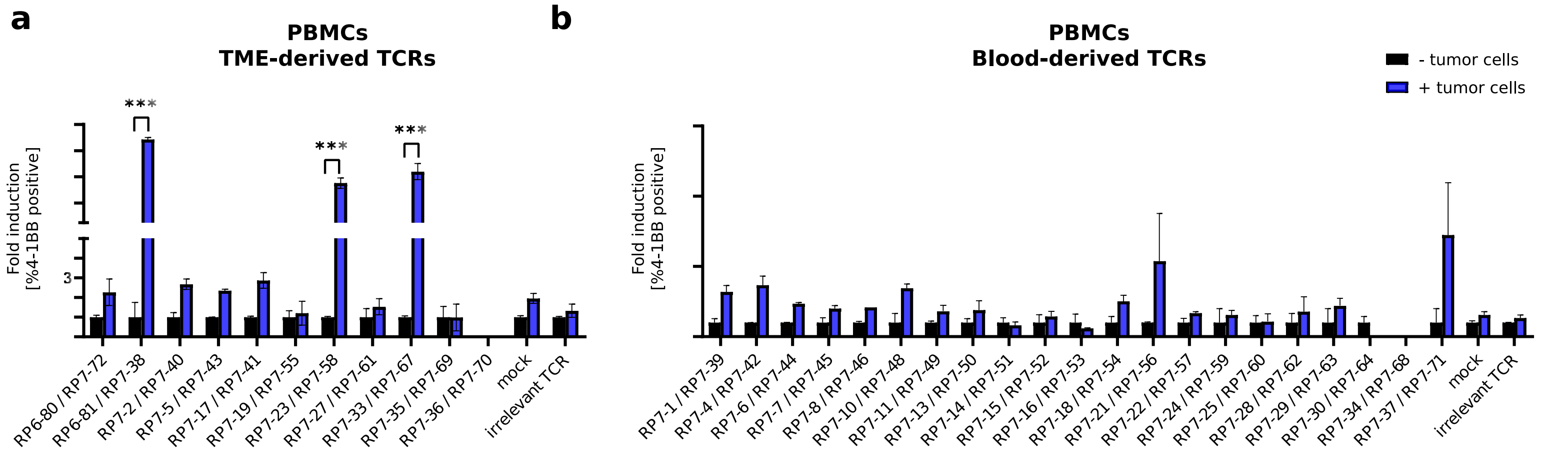


**Extended Data Figure 6.**
